# Coevolution of phenological traits shapes plant-pollinator coexistence

**DOI:** 10.1101/2024.08.08.607160

**Authors:** François Duchenne, Virginia Dominguez-García, Francisco P. Molina, Ignasi Bartomeus

**Affiliations:** Estación Biológica de Doñana (EBD-CSIC), Sevilla, Spain; Swiss Federal Institute for Forest, Snow and Landscape Research (WSL), 8903 Birmensdorf, Switzerland

**Keywords:** feasibility, emergent properties, morphology, structure, competition, plant, pollinator

## Abstract

Previous research has revealed how species traits determine species interactions, and how species interactions influence species coexistence. However, this hierarchical view ignores the coevolutionary feedback from species interactions to species traits and its consequences for species coexistence. Here, we developed a theoretical model of quantitative genetics to explore how the coevolution of morphological and phenological traits shapes the structure and stability of mutualistic interaction networks. We found that, in comparison to morphological traits, phenological traits led to distinctive species evolutionary trajectories, resulting in different emergent properties at community level. This is because phenological traits promoted facilitation over competition. While morphological coevolution gave rise to modular and specialized interaction networks with a negative diversity-stability trade-off, phenological coevolution was required for the emergence of nested interaction networks that exhibited a positive relationship between diversity and structural stability. Empirical observations from 17 pollination networks were consistent with the theoretical results: we found many phenological motifs promoting facilitation over competition, suggesting an important role of phenological coevolution in community assembly. The seasonal organization of empirical interactions enhanced the community stability and dampened the diversity-stability trade-off that was observed when aggregating interactions across time. Our results highlight the importance of phenological coevolution in the emergence of diverse and stable communities.

## Introduction

Over the last decades, the study of mutualistic communities has clarified the ecological determinants of their stability, advancing our understanding of how diverse communities can be ecologically stable (1–8). In mutualistic systems, species persistence depends on the balance between three important kinds of effects that each species receives: the positive effects generated by direct mutualistic interactions between guilds, the negative effects generated by direct competitive interactions among species of the same guild, and the indirect effects propagating through chains of species interactions, which can be positive or negative. The balance between these three effects is strongly affected by the way the species interactions are distributed in the community (1, 2, 4). For example, nestedness, a structural pattern which occurs when specialist species interact with generalist ones, has been shown to increase community persistence by promoting positive indirect effects (facilitation), which can overcome the negative effects of direct competition (1, 4, 9). More recently, several studies have suggested that the temporal distribution of the interactions, a previously neglected facet of network structure, is also key for understanding the competition-facilitation balance and the stability of mutualistic communities (5, 10, 11).

The growing availability of temporally resolved mutualistic networks has revealed strong seasonal dynamics, with high turnover in species and interactions across seasons (12–14). These temporal dynamics are partially determined by species phenologies (*e*.*g*. flowering period, activity period; 12, 14–16). This species turnover can decouple interactions between different partners in time, and thus, decouple the mutualistic interactions between guilds and the within-guild competition for partners. This decoupling generates positive indirect effects, *i*.*e*. facilitation (5, 10, 11). For example, in a plant-pollinator community, two pollinators can interact with the same plant population but at different periods of the season: an early pollinator may do so in spring and a later one in midsummer. This leads to facilitation between the two pollinators, because they maintain the same plant population, without direct competition. Since they promote positive indirect effects, phenological traits are more likely than morphological traits to prevent competitive exclusion of specialist species (*i*.*e*. species with few partners) (10). These specialist species are the most vulnerable species but also the ones contributing the most to the nested structure of mutualistic networks (17), which promotes community stability (1). Thus, phenological traits have been shown to be key for the ecological stability of mutualistic networks, but all these studies have been done with fixed trait values, without considering the evolutionary processes shaping species traits.

When considering evolutionary dynamics, we expect phenological coevolution to generate more nested and stable mutualistic networks than morphological coevolution. This is because phenological traits can enhance positive indirect interactions, fostering a broader spectrum of species’ generalism – from specialists to generalists – which is required to produce nested networks. However, to have a positive effect on the balance between facilitation and competition, phenologies need to be organized so that mutualistic partners overlap in time, while competitors do not (Fig. 1). Thus, one of the main questions that remains is how easily these patterns can emerge from coevolutionary dynamics in species-rich communities? Since natural selection is driven by differences in fitness and not by community properties, it is not obvious that evolutionary dynamics would lead to a network structure promoting community stability.

**Fig. 1:**
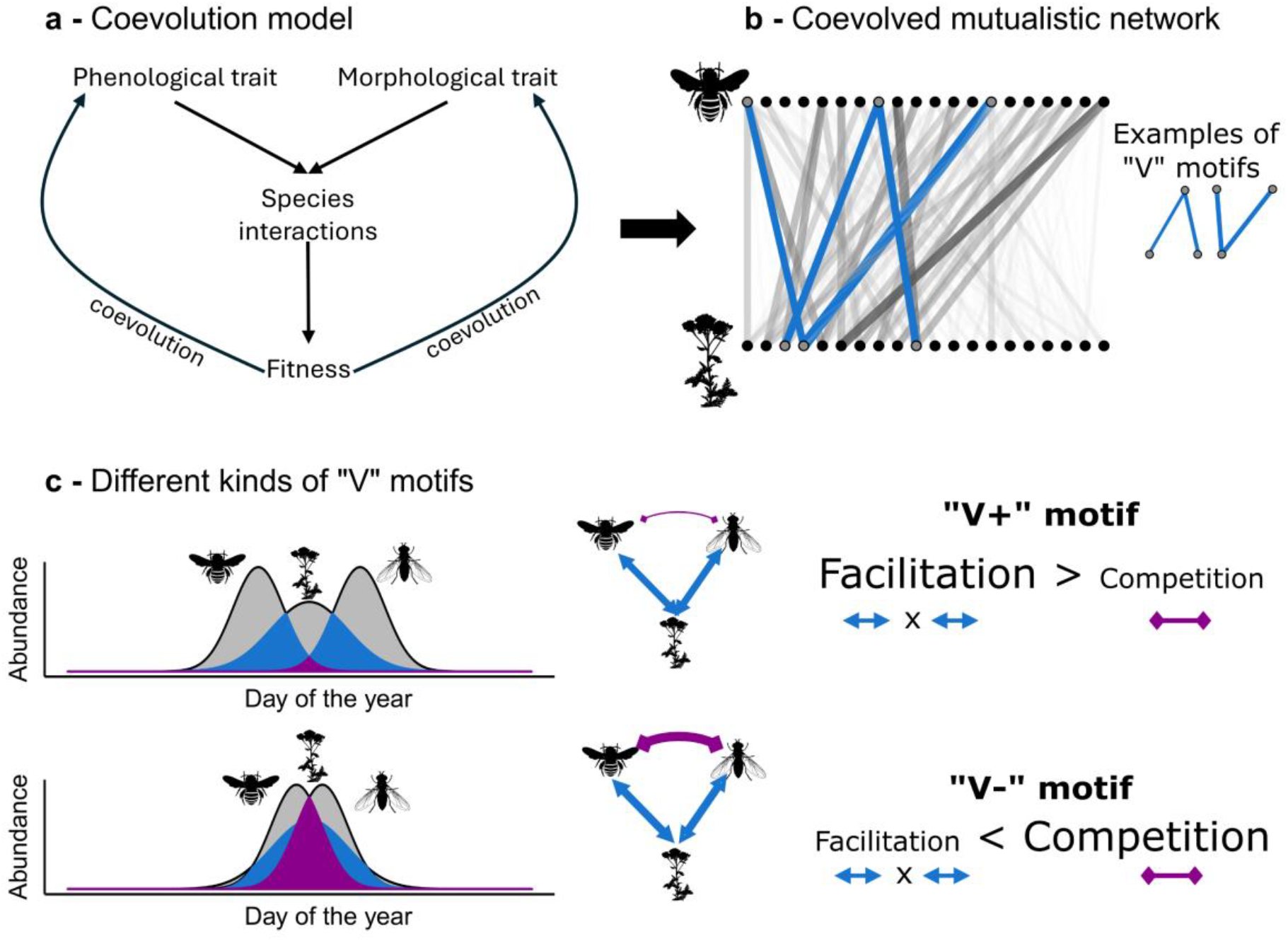
schematical representation of our coevolution model and the different motifs that can emerge from phenological coevolution. (a) Schematical representation of our theoretical model and (b) of a coevolved mutualistic interaction network produced by that model. Each dot is a species of pollinator (top row) or plan (bottom row). Line thickness is proportional to interaction strength. Interaction networks contain many “V” motifs of 3 species, such as the two highlighted in blue. These motifs can be classify as “V+” or “V-” motifs depending on whether they promote facilitation over competition or not. (c) On the top a “V” motif promoting facilitation (“V+”), and on the bottom a V motif promoting competition (“V-”). On the top schema, pollinators interact with the same plant but overlap little in time, leading to weak interaction strengths but very low competition relative to facilitation. The facilitation between pollinators is the positive indirect effect mediated by the plant (the product of the mutualistic interactions). On the bottom schema, pollinators largely overlap in time, a structure which tends to promote strong interaction strength and stronger competition than facilitation, which can lead to competitive exclusion. For simplicity we focused on facilitation and competition between two pollinators, but opposite “V” motifs, with two plants and one pollinator, follow the same logic.

Answering that question requires addressing some methodological issues that limit our understanding of how coevolution shapes the stability of mutualistic communities. First, the few studies that have explored coevolution in mutualistic communities have focused, explicitly or implicitly on the coevolution of morphological traits (18–24), neglecting the temporal organization of interaction networks. Second, these studies have also used different metrics of stability, using either the proportion of species persisting (19, 20, 24), the resilience (18, 22, 23) or the robustness (21) to a perturbation, which complicates their synthesis. Third, most of the coevolutionary models applied to mutualistic systems have neglected the competition dynamics that are coupled with changes in mutualistic interactions, but see refs. (20, 22) for exceptions. Understanding how coevolution shapes the stability of mutualistic systems requires modelling the coevolution of morphological and phenological traits within the same framework, because both have been shown to determine mutualistic and competitive interactions in nature (14, 15, 25). To overcome these limits, we thus need to develop modelling frameworks that allow us to: (i) include the coevolutionary dynamics of phenological traits in addition to morphological ones, (ii) use a stability metric that is easily comparable across simulations, (iii) account for competition and mutualism dynamics.

To investigate the potential emergence of a stabilizing structure in mutualistic communities due to the coevolution of phenological and morphological traits, we first developed a theoretical model of quantitative genetics to track interaction network structure over the coevolution process. We then coupled that model with a structural stability approach to compare the stability of networks produced by coevolution of morphological and/or phenological traits. Finally, we analyzed a set of 17 time-structured empirical plant-pollinator networks to compare the observed seasonal organization of mutualistic interactions (henceforth seasonal structure) with those produced by our coevolution model. Our results show that a seasonal structure reducing competition and stabilizing communities can emerge from the coevolution of phenological traits, and that this kind of structure can be found in empirical systems. This suggests that the study of temporal dynamics in community ecology can help to answer the longstanding question of how diverse natural communities remain stable.

## Methods

We developed a theoretical model of quantitative genetics to simulate the coevolution of phenological and/or morphological traits of two guilds of species interacting mutualistically between guilds and competing within guilds (Fig. 1). In this model morphological and phenological traits determine direct species interactions: mutualistic interactions among guilds and competitive interactions within guilds (Fig. 1a). Species interactions determine population fitness, which drives trait evolution, leading to coevolution among species (Fig. 1a). Our model allowed us to track, as the coevolution dynamics unfolded, the structural properties of the interaction network and the probability that all species coexist, given the interaction network (*i*.*e*. its structural stability). To assess the role of phenological and morphological coevolution in determining network structure and stability, we performed simulations over three different scenarios: with a phenological trait, with a morphological trait and with both traits together.

Here, for the purpose of describing the model we refer to a plant-pollinator community, but the model could apply to any mutualistic community. Below, we develop all the equations for the full model (with phenological and morphological traits) and for the plant side, but pollinators were treated in a similar way (equations are shown in Supplementary Methods I).

### Species traits

Each species, consisting of a single population, was characterized by two Gaussian traits: a morphological trait and a phenological trait, with a mean *μ*_*M*_ and *μ*_*P*_ and a standard deviation *σ*_*M*_ and *σ*_*P*_, respectively. To make comparisons possible between traits, both were treated exactly in the same way, on a scale that goes from 0 to 365.

For phenology, the trait distribution reflects the day of the year and thus models the entire duration of the activity and flowering periods of pollinator and plant populations, respectively. Similarly, the morphological trait can be view as a projection onto one dimension with an arbitrary scale, of a niche based on two traits. For example, the mean parameter could be view as the proboscis or corolla length, of pollinators and plants, respectively, while the standard deviation would be defined by another trait mediating flexibility in morphological matching, such as the width of the pollinator proboscis and the opening width of the flower corolla.

To maintain gaussian distributions within the interval [0,365], we bounded the mean trait value and standard deviation between 80 and 285 and between 1 and 50, respectively. These constraints on trait values had advantage to avoid infinite evolution but also mimicked physiological limits. For example, for phenology, these limits constrained pollinators to be active mostly between days of the year 80 and 285 (*i*.*e*. spring and summer) simulating empirical pollination seasons that we can observe in temperate areas. For morphology, it would correspond for example to a limit in the length of the corolla or feeding apparatuses, that cannot be too short or too long because of physiological constraints. To bound trait values without using a sharp truncation that would create artefacts at the bounds, we used a process inspired by generalized linear models, in which the trait evolves on an unbounded latent distribution linked to the effective bounded values through a link function (*cf*. Supplementary Methods I). In the following part, *μ* and *σ*, denote the values on the bounded scale (real mean and standard deviation), while 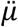 and 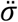 denote the value on the latent unbounded scale. For example, 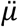 varied from −∞ to +∞, but *μ* was bounded between 80 and 285 (Fig. S1).

### Species interactions

The overlap between their morphological (*φ*_*ij*_) and phenological traits (*τ*_*ij*_) determined the per-capita interaction strength (*γ*_*ij*_) between a pollinator *j* and a plant *i*:

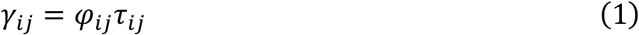

To calculate the overlap between traits, we used the area below the minimum of the two gaussian traits involved (Fig. 1c):

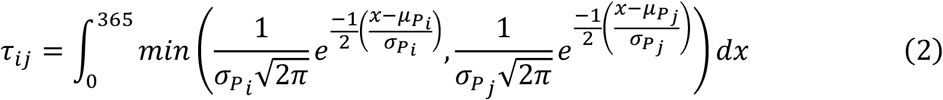

*φ*_*ij*_ was calculated according to an equation similar to equation (2). Equation (1) and (2) means that the per-capita mutualistic interaction strengths are bounded between zero and one (0≤*γ*_*ij*_≤1), because *φ*_*ij*_ and *τ*_*ij*_ are bounded between zero and one. To neglect phenology in simulation with only a morphological trait, we set τ =1 for all pairs of species, meaning that all interactions occur simultaneously. In contrast, to ignore morphology when simulating with a phenological trait only, we set φ =1, meaning that all species pairs have a perfect morphological match.

The competition between plant *i* and *k* was defined as the intersection of their mutualistic interactions divided by the union of their niche, multiplied by the temporal co-occurrence between plant *i* and *k* (*τ*_*ik*_):

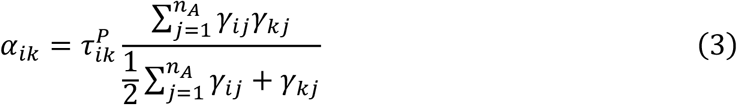

where *n*_*A*_ was the number of pollinator species and 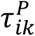 the within guild phenological overlap between plant *i* and *k*.

The competition coefficient among pollinator species was calculated with a similar equation (*cf*. Supplementary Methods I). For plants and pollinators, we set the intraspecific competition to one, *α*_*i,k*=*i*_ =*α*_*j,k*=*j*_ =1, to ensure diagonal dominance of our system. These competition coefficients represent only the competition for mutualistic partners among species from the same guild, as here we ignored competition for abiotic resources or space, because we wanted to focus on the role of mutualistic interactions in driving coevolution.

### Species fitness and trait evolution

The fitness of each population was defined as the balance between competition and mutualism that each population received. For example fitness of plant *i* (*W*_*i*_) was defined as follow:

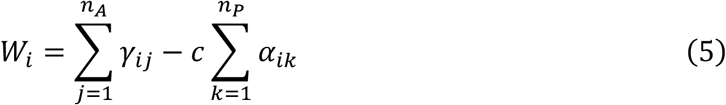

The fitness function of a pollinator *j* (*W*_*j*_) was calculated with a similar equation (*cf*. Supplementary Methods I).

At each time step, each population trait parameters evolved according to the breeder’s equation (26, 27):

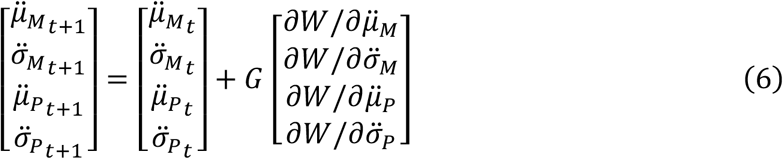

where G is the additive genetic variance-covariance matrix. For simplicity, we assumed that trait parameters were independent and that they all had the same additive genetic variance (*g*). Thus, *G* was a 4×4 diagonal matrix:

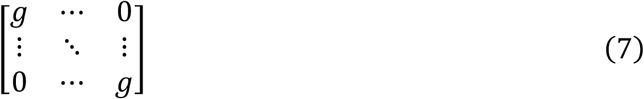

Although morphological and phenological traits evolved independently at each time step due to selection, when they were included in the same simulation, their evolutionary trajectories were not independent, as they affected the same elements of fitness, namely mutualistic and competitive interactions.

Since selection was implemented on the latent unbounded scale of the trait parameters, evolution speed asymptotically slowed down to zero when reaching the bounds of the trait space, thus maintaining trait value in the defined trait space (Fig. S1).

### Simulations and parametrization

Our simulations required very few parameterizations, because most of the coefficients were determined by the evolving trait values, but we needed to draw initial values for traits and to fix a value for the competition importance (*c*) and for the additive genetic variance (*g*).

For the traits, we drew initial values on the modified logit scale (*cf*. Supplementary Methods I) from a uniform distribution for the mean and the standard deviation: 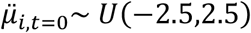 and 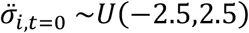. We generated 100 initial conditions for 3 different values of initial diversity: 10, 20 and 30 species per guild.

Since our competition coefficients were always smaller than the per-capita mutualistic interaction strength (*cf*. equation (2)), we needed *c*>1 to avoid mutualistic orgies, a common behavior in linear models of mutualistic communities (28). A mutualistic orgy would lead to all species quickly converging to the same trait value, and would thus not be realistic. We performed simulations with three values of competition to simulate 3 cases: low importance of competition (*c* =2), a moderate importance (*c* =4) and a high importance (*c* =6). These three values were chosen to have an average ratio of mutualism/competition across the 100 initial conditions (when considering a phenological trait) that was >1 (mutualism stronger than competition), 0.5<x<1 (competition slightly stronger than mutualism) and <0.5 (competition stronger than mutualism, Fig. S2).

The additive genetic variance *g* was fixed to 0.01, which allows a smooth evolution but fast enough to allow simulation to reach a stationary state (Fig. S3).

We performed simulations over 5000 time steps. In our time-discrete model, trait values never reach an equilibrium because all species traits change simultaneously, but trait evolution was fast in the first 500 steps, before slowing down and reaching stationary states asymptotically (Fig. S3). For each initial condition, the simulation was run three times, considering a morphological trait or a phenological trait only, and considering both traits together. This allowed us comparing trait coevolutionary dynamics in isolation and when they were mixed.

### Network structure, motifs and stability

To assess how coevolutionary dynamics affect the structure and the stability of the evolving interaction networks, we measured indices describing the mutualistic interaction network. Since evolution slowed down with time (Fig. S3), we measured these metrics at 10 time steps, that were evenly distributed on a square-rooted time scale (t = 0, 60, 250, 560, 990, 1540, 2220, 3020, 13950, 5000).

To measure how evolution affected the balance between mutualism and competition we computed the average competition strength, which is the average of all *α*_*ij*_, and the balance between mutualism and competition as 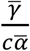, where 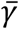and 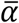are the average of all pairwise terms of mutualistic interaction and competition (across both guilds), respectively. To test if co-evolutionary dynamics can lead to phenological motifs that promote facilitation over competition, among the hundreds or thousands of “V” motifs we calculate the proportion of them promoting facilitation, *i*.*e*. “V+” motifs (Fig. 1c). This kind of motifs can be present only in the phenological scenario because they require that two mutualistic partners of a given species are decoupled in time. In contrast, when considering morphological traits, all interactions happen simultaneously (*τ* =1) thus “V+” motifs are impossible. Details of the calculation are provided in Supplementary Methods II.

To measure structural properties we measured the connectance, interaction evenness, specialization, nestedness and modularity with different indices implemented in the *bipartite* R package (29). These structural properties measured different, but not independent, facets of the overall network structure. Since we worked with quantitative networks, here network connectance was not measured as the proportion of realized interactions but as the average diversity of partners across species. This connectance can be expected to be positively related with interaction evenness, which measures how heterogeneous interactions strengths are. To complement these metrics, we also included in these analyses the average mutualism strength 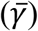which measure how full is the mutualistic interaction matrix and thus can be view as another measure of weighted connectance. At contrary, specialization measures the average tendency of species to interact with specific partners. Finally, nestedness and modularity described the possibility to arrange the matrix to have most of the interactions clustered in the upper triangle of the interaction matrix or in several independent blocks of species, respectively. Details of the calculations are provided in Supplementary Methods II. To track network structure over time, we performed a Principal Component Analysis (PCA) with the structural metrics, across all simulations and time steps. We used the two first dimensions of this PCA which explained 93% of the total variance, to characterize the evolutionary trajectories of the simulated communities.

Finally, to assess the stability of the interaction networks produced by coevolutionary dynamics, we measured the community structural stability. Structural stability is a measure of ecological stability that aims to quantify the likelihood of permanent coexistence of all species, according to their interactions (2, 3, 30). To compute this coexistence likelihood, we used a generalized Lotka-Volterra in which species interactions are extracted from the networks of competitive and mutualistic interactions produced by coevolution dynamics (Supplementary Methods II). Population dynamics of a given species was thus defined by its interactions with other species and by a per-capita growth rate (*r*), which defines how that species grow in isolation, for a given environmental condition.. The range of environmental conditions, or growth rates, in which all species can coexist is called the feasibility domain. The structural stability is quantified as the size of the feasibility domain scaled by the number of species in the community (*ω*), to compare that metric across communities with different number of species. This stability metrics has the advantages of not depending on an arbitrary parametrization of growth rates or initial abundances, but to depend only on the network of species interactions. To calculate the structural stability of our communities, we used the *Omega* R function provided by Song *et al*. (30). Details of the calculation are also provided in Supplementary Methods II.

### Diversity-Stability relationship

To study how coevolution affected the relationships between stability and diversity (i.e. species richness), we modelled structural stability as a function of the number of species per guild using a Generalized Linear Model (GLM). This GLM had a quasibinomial error structure and a logit link function. To model one diversity-stability relationship per coevolution scenario, for initial and coevolved networks, we included a quadruple interaction between the diversity (number of species per guild), the kind of coevolution scenario, the kind of network (initial *versus* coevolved) and the importance of competition in the model. The GLM was described by the following equation:

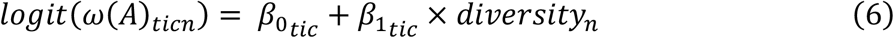

where *ω*(*A*)_*ticj*_ was the structural stability at time *t*, of the initial (*t* = 0) or coevolved networks (*t* = 5000), for coevolution scenario *i* (phenological, morphological or mixed coevolution), competition importance *c*, and for a simulated diversity *n* (number of species per guild). 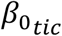and 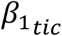were the intercept and the slope of the diversity-stability relationships, respectively.

### Analyses of empirical plant-pollinator interaction networks

To assess the contribution of seasonal structure of natural mutualistic communities to their stability in nature, we used a set of 17 time-structured plant-pollinator interaction networks, sampled in the Doñana National Park, an area near Sevilla (Spain) with a mediterranean climate. Pollinators were insects only, mostly bees (Hymenoptera, Apidae) and Hoverflies (Diptera, Syrphidae), while flowering plants were mostly mediterranean shrubs: *Cistus, Lavandula* and *Salvia* being the most common genus. These networks have been sampled at bi-weekly intervals during the pollination season (February to May), for one to 8 years, with the protocol described in *ref*. (31). Interaction networks include between 21 and 131 insect species and between 8 and 36 plant species (Table S1). Interactions have been recorded at each sampling occasion along a 100m transect during 30 effective minutes, and flower abundances per species have also been recorded along that transect in a standardized way (31).

From the empirical data, we could extract species phenologies and temporal information about the occurrence of pairwise interactions. To assess how the seasonal structure in interactions affected the structural stability of the empirical communities, we estimated the structural stability (*ω*) in two ways: we either aggregated over time (i.e. removing seasonal structure) or kept the seasonal structure in interactions, thus using the empirical phenological overlaps (*cf*. Supplementary Methods III). Although we did not have information about morphological traits, we assumed that by using empirically observed interactions in both scenarios (aggregated or not), the species interactions already depended on the putative morphological match between plants and pollinators. Thus, we could estimate the effect of the seasonal structure on network stability, accounting for other factors determining species interactions (morphology, etc.).

Additionally, we compared the proportion of “V+” motifs found in empirical communities, with the ones predicted by simulations, using either coevolved or randomly distributed species traits. From the empirical interaction data and flower counts we could assess the number of “V+” motifs, in which the (geometric) average overlap between mutualistic partners was higher than the overlap between the two competitors. To get a measure comparable with our simulations and among networks, as before, we estimated the proportion of “V+” motifs over the total number of “V” shape motifs (*cf*. Supplementary Methods III). To compare these proportion of “V+” motifs with the proportion we obtained in our theoretical simulations, we performed a new set of simulation using the number of plant and pollinator species of the empirical networks. For each of the 17 empirical networks, we performed these simulations with a mixed coevolution scenario (morphological and phenological coevolution together), for a moderate importance of competition (*c* = 4) and for a set of 10 different random initial conditions of randomly distributed traits values, using the same uniform distribution as for the theoretical simulations to draw trait values.

## Results

The initial average competition strength differed between initial networks with or without phenology (Fig. 2), because decoupling interactions in time, even randomly, decreases competition strength. Moreover, the scenario combining both phenological and morphological traits exhibited less mutualism strength in average at initial state, because combining two randomly distributed traits provided less opportunity for trait matching than when using only one trait. However, what most interested us was seeing how coevolution shaped the networks, *i*.*e*. the trajectory over time of the different structural metrics. When coevolutionary dynamics unfolded, trait modifications quickly reorganized species interactions, affecting the overall strength of competitive and mutualistic interactions and their balance, with distinctive trajectories across coevolution scenarios (Fig. 2 & S4).

**Fig. 2:**
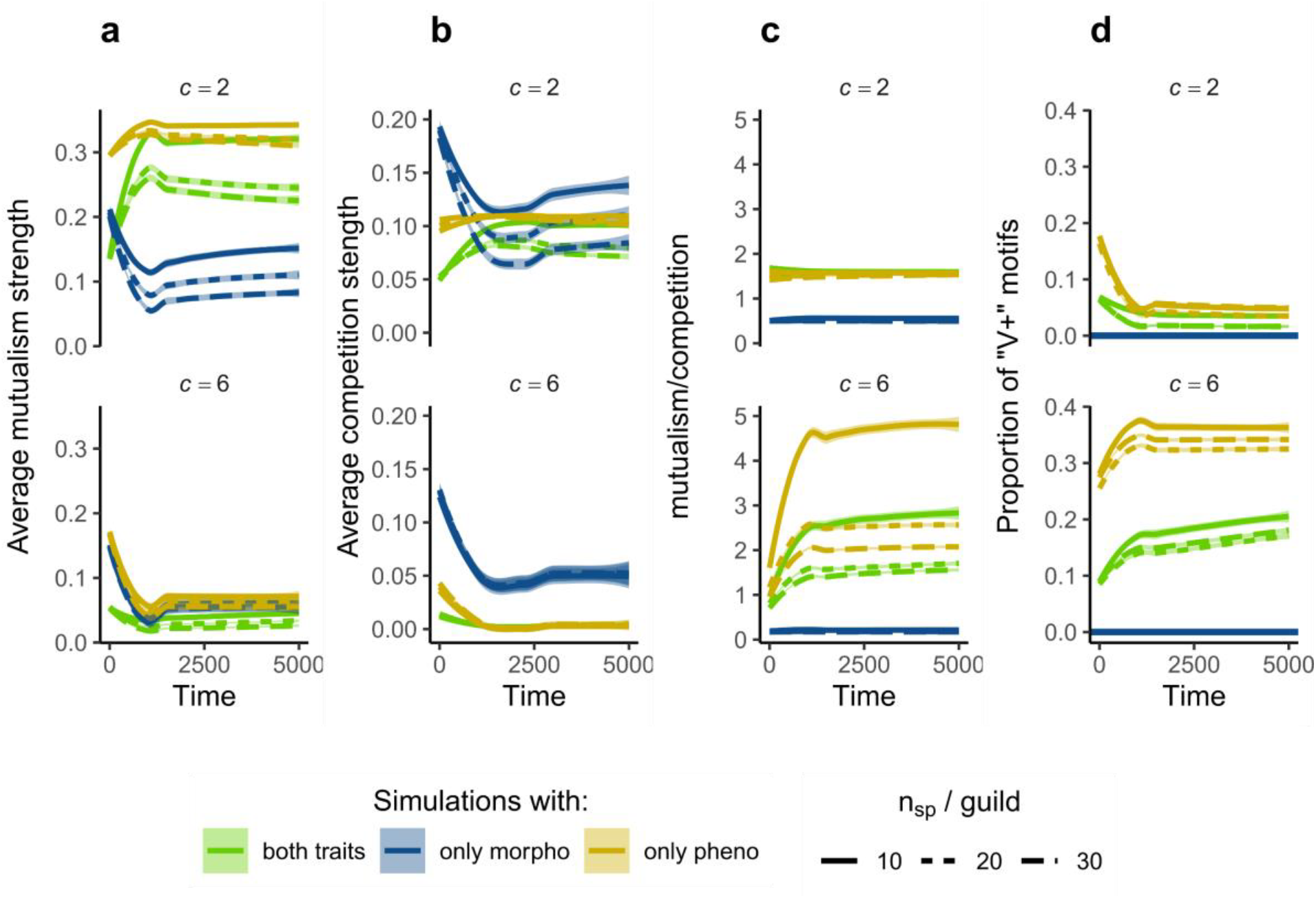
motifs promoting facilitation can emerge from phenological coevolution. (a) average mutualistic strength, (b) average competition strength, (c) mutualism/competition ratio (cf. methods) and (d) proportion of “V+” motifs (over all “V” motifs), as a function of time, diversity and simulation scenarios for low (c=2) and high (c=6) competition strengths. In (a)-(d) the lines and ribbon show the estimated values and their 95% confidence interval, respectively, obtained by a locally estimated scatterplot smoothing (LOESS) of the values across time. V+ motifs did not exist when performing coevolution with morphological trait only (cf. Methods). See Fig. S4 for simulations with c = 4.

Regardless the importance of competition (parameter *c*), morphological coevolution reduced the average level of mutualistic benefits and consequently also reduced the average level of competition for partners (Fig 2a-c). Since competition decreased more strongly than mutualism benefits, morphological coevolution slightly increased the mutualism-to-competition ratio; however, this ratio remained below one, meaning that competitive effects were still stronger than mutualistic benefits. This restructuring of the community can be understood by examining the distribution of species traits: morphological coevolution produced only specialist species, characterized by the small standard deviation of their morphological trait, indicating a very narrow morphological niche (Fig. S5).

In contrast, phenological and mixed coevolution produced some generalist species, characterized by long phenology (large standard deviation, Fig. S5), in a proportion that depended on the importance of competition. The lower the competition importance, the more generalist species (Fig. S5), and thus the more mutualism and competition in the community (Fig. 2a-b). When giving low importance to competition (*c* = 2), phenological and mixed coevolution tended to maximize trait matching across a maximum of species, leading to trait convergence, regardless the amount of competition it generated. In contrast, when competition had moderate or high importance (*c* = 4 or *c* = 6), phenological and mixed coevolution produced interaction networks that tended to maximize the mutualism-to-competition ratio (Fig. 2c & S4). Under these conditions, phenologies were reorganized in a way that potential competitors did not overlap in time. This reorganization creates “V+” motifs that promote facilitation (Fig. 2d & S4), patterns that cannot be produced by morphological coevolution.

To understand how coevolution affected network structure and stability, we then focus on simulations in which competition was moderate (*c* = 4) or important (*c* = 6), as it appears to be the case in many mutualistic systems (32–34). In this context, our simulations highlighted significant differences across various aspects of network structure between the coevolution scenarios including phenological traits (mixed or phenological coevolution) and the scenario with morphological traits only (Fig. 3 & S6). Mixed and phenological coevolution tended to produce mutualistic networks with a very similar structure, more nested and less modular than mutualistic networks produced by morphological coevolution. Differences in network structure between coevolution scenarios became apparent mostly when simulating large networks (*i*.*e*. high number of species, Fig. 3d-e), which offers a wider range of possible interactions and thus more degree of freedom for the emergence of structural patterns.

**Fig. 3:**
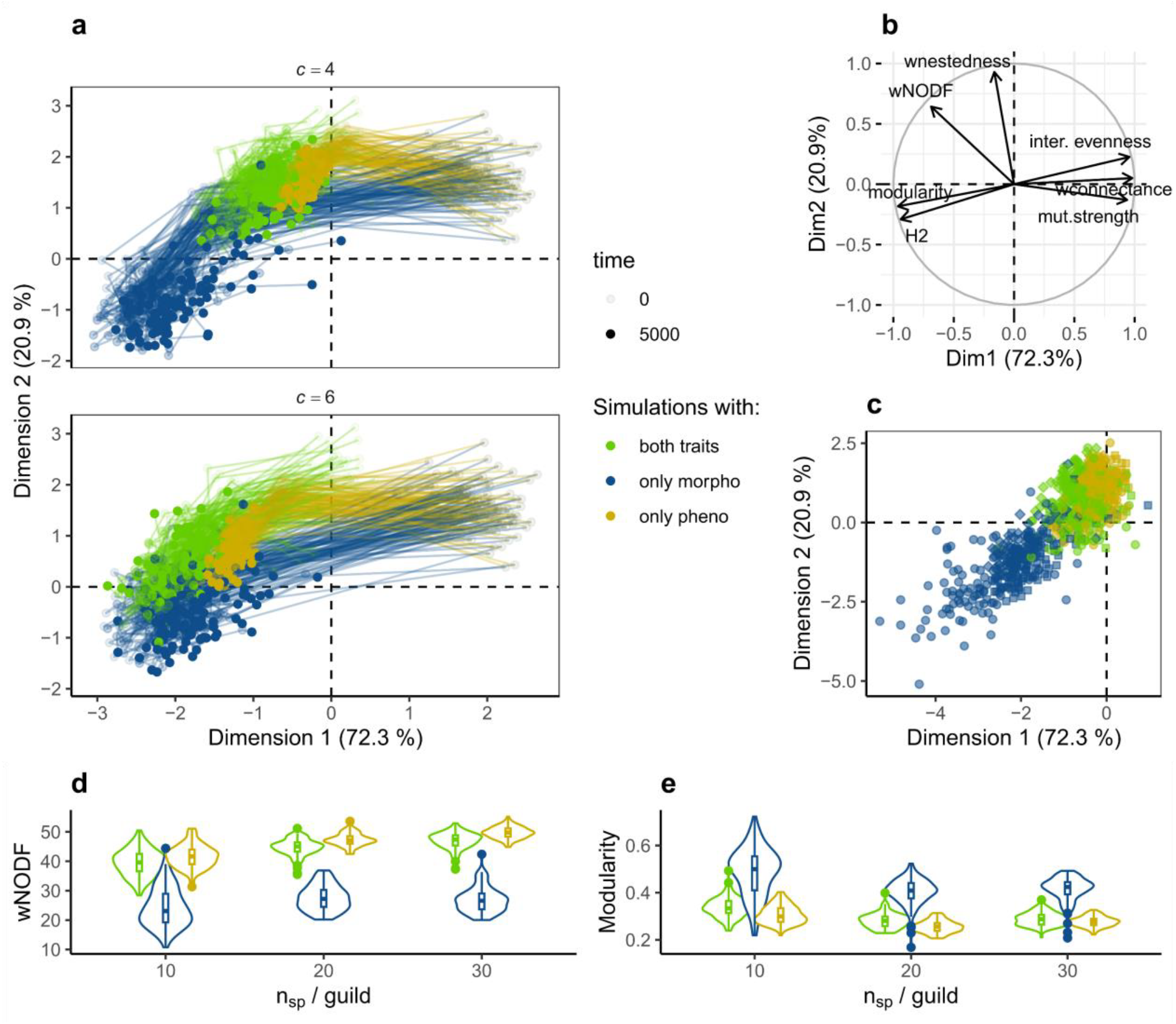
phenological and morphological coevolution led to divergent evolutionary trajectories of communities. (a) Location of each network at different time steps, in a multidimensional space of network structure, for different importance of competition (c = 4 & c = 6). The multidimensional space of network structure was created using the Principal Component Analysis (PCA) illustrated in (b). In (a) only the simulations with 30 species per guild are represented for readability. (b) PCA loading plot, where wNODF is a weighted measure of nestedness and H2 a measure of specialisation, and mutualism (mut.) strength a measure of connectance (cf. methods). (c) Final location of each community in the multidimensional space of network structure, when c = 4 (cf. Figure S6 for c = 6). Circles represent simulations with 10 species per guild, squares 20 species per guild and diamonds 30 species per guild. (d) and (e) represent the nestedness (measured as wNODF) and the modularity of the final networks, as a function of the number of species per guild and kind of trait, when c =4.

These first results suggest that phenological coevolution is more likely to decrease the competitive exclusion than morphological coevolution, because of a better optimization of the ratio between mutualism and competition (Fig. 2), and because of the emergence of stabilizing structural properties, such as nestedness (Fig. 3). Importantly, when both traits coevolved, the outputs of the coevolved networks were more similar to the ones obtained by phenological coevolution than to the ones obtained by morphological coevolution, suggesting that coevolution of species phenologies is key in determining emergent properties at community level.

In the initial state, communities structured by morphological traits exhibited lower structural stability than those structured by phenological traits or by both traits (Fig. 4a). This is consistent with previous findings (5, 10), and happens because decoupling interactions in time, even randomly, decreases competition strength (Fig. 2b-c). All initial networks showed a negative relationship between diversity and stability: the more diverse, the less stable (Fig. 4b-c). Once the coevolutionary dynamics unfolded, they increased the structural stability and dampened the diversity-stability trade-off in all coevolution scenarios (Fig. 4b-c). However, only the phenological and mixed coevolution reversed the diversity-stability trade-off to a positive relationship (Fig. 4b-c). This suggests that the seasonal structure in interactions is key to create diverse and stable communities.

**Fig. 4:**
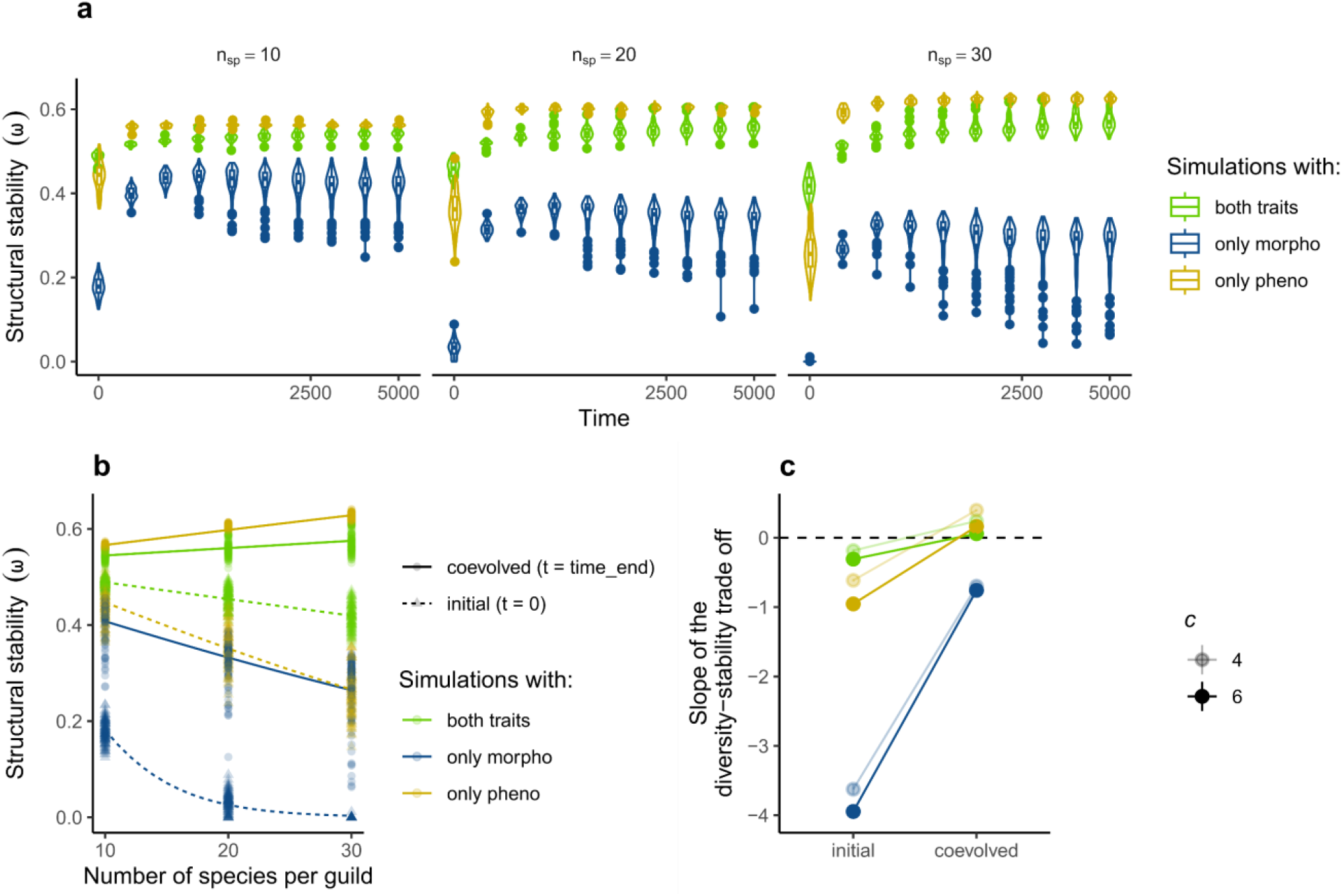
phenological coevolution increased structural stability and dampened the diversity-stability trade-off. Changes in structural stability (average species persistent probability) over time, for the three coevolution scenarios and number of species per guild, when competition was moderate (c = 4). See Fig. S7 for simulations with c = 2 & c=6. The x-axis is square-root transformed. (b) structural stability as a function of the number of species per guild (i.e. diversity), for interspecific competition strength c=4. Lines show the predictions of a Generalized Linear Model (GLM, cf. Methods). (c) Slopes of the diversity-stability relationships extracted from the GLM. Error bars (±CI_95%_) are too small to be visible.

Finally, to assess the stabilizing role of phenology on real-world communities, we analyzed the phenological organization of 17 empirical plant-pollinator networks. We found evidence for a high proportion of the “V+” motifs (Fig. 5a), described in Fig. 1c, which suggests that coevolution of species phenologies have shaped the seasonal structure of plant-pollinator interactions. Our results show that the proportion of motifs promoting facilitation were relatively consistent with what was predicted by mixed coevolution and higher than what would be found in random initial networks (Fig. 5a).

**Fig. 5:**
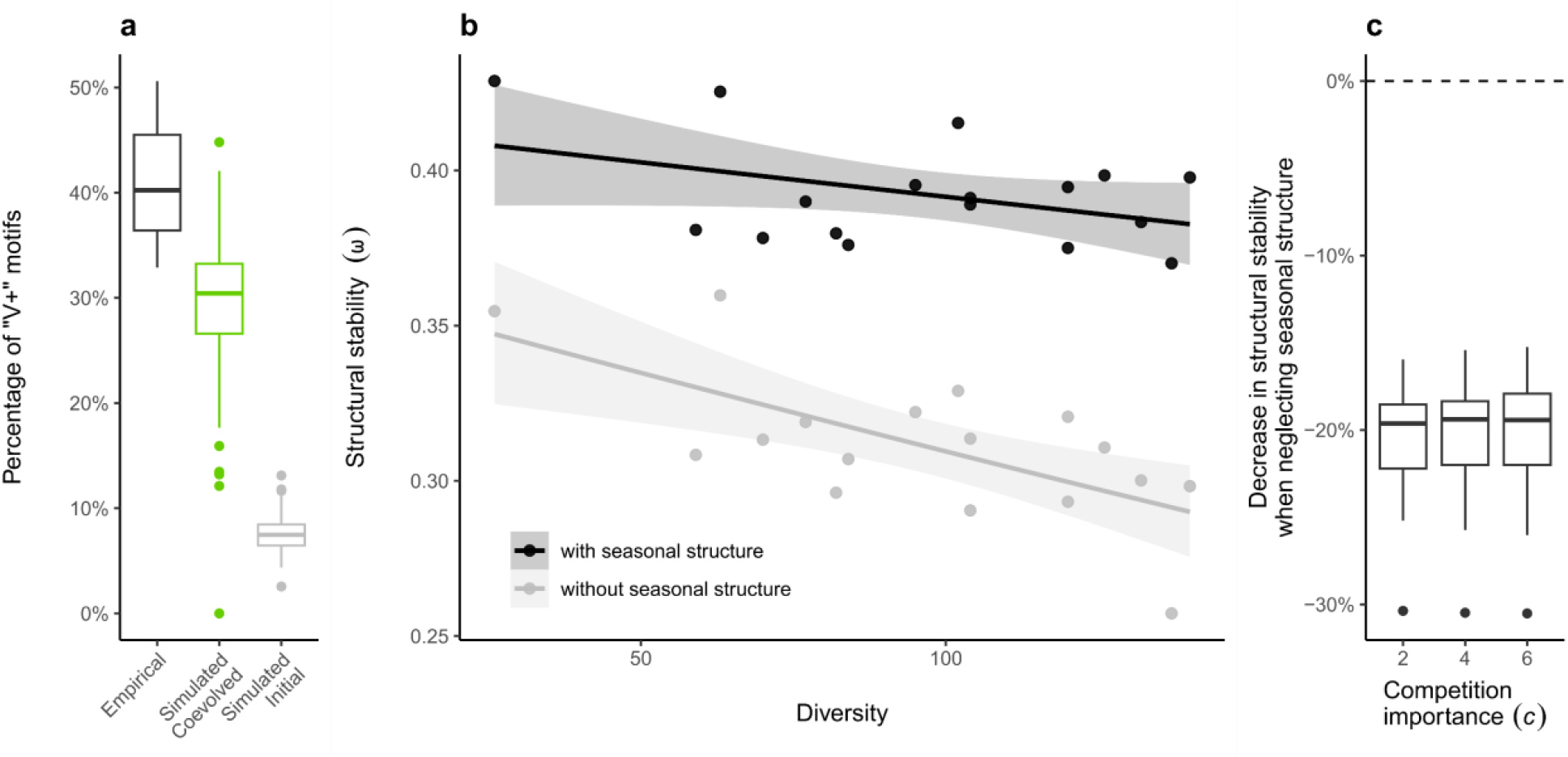
Seasonal structure in interactions stabilize natural communities. (a) Distribution of the percentage of “V+” motifs, relative to the total number of V motifs, in the 17 empirical networks, and in simulated networks with similar dimensions (cf. methods), before (i.e. random, grey) and after mixed coevolution (green). (b) Structural stability, average probability of persistence, as a function of community diversity (number of pollinator and plant species), when accounting for seasonal structure in competitive interactions or when neglecting this seasonal structure, for a given competition importance (c = 4). The lines and ribbon show the estimated values and their 95% confidence interval, respectively, obtained by generalized linear model with a quasi-binomial error structure. (c) The decrease in percentage of the structural stability observed when neglecting seasonal structure, relative to the measure when considering it, as a function of the overall competition importance (c parameter). n = 17 for each boxplot. Dots represent outliers.

As predicted by the simulations, considering the seasonal structure in interactions led to an increase in the structural stability of the empirical communities (Fig. 5b-c). This shows that the stabilizing role of the coevolved seasonal structure that we found *in silico* is likely to be important in stabilizing natural mutualistic communities. In addition, the negative relationship between stability and diversity that we observed in empirical communities was dampened when seasonal structure was considered (Fig. 5b).

## Discussion

Our results show that explicitly modelling the linkage rules defining species interactions is important to understand the community level outcomes of coevolutionary dynamics. We show that, in theory, a positive relationship between diversity and stability could be achieved by the coevolution of phenological traits but not by the coevolution of morphological traits. Consistently with our theoretical results, we show that the seasonal structure of empirical plant-pollinator communities strongly dampened the diversity-stability trade-off observed when neglecting that seasonal structure. Our results highlight that the temporal organization of species interactions, which emerges when phenological traits are considered, is key to understand coevolutionary dynamics and the stability of diverse mutualistic communities.

In contrast to morphological traits, phenological traits can decouple interactions in time, rendering an asymmetric partition between the mutualistic interactions and the competition for mutualistic partners (10, 35). Surprisingly, this key mechanism explaining species coexistence has only recently been integrated to ecological models, but is now well recognized as an important ecological mechanism (5, 10, 11, 35–39). However, most of these studies consider that species phenologies are mostly determined by abiotic factors (10, 37, 39). Here we show that a seasonal structure fostering stability can emerge from coevolutionary dynamics only. Our model did not include any abiotic constrain on phenologies other than physiological bounds, common to all species. Thus, in our simulations, the emergence of the seasonal structure was only driven by species interactions. This suggests that the mechanism we describe could be universal: a seasonal structure in interactions is expected to be present in every mutualistic system where interspecific competition for mutualistic partners is high enough, regardless the abiotic seasonality. Since, interspecific competition for mutualistic partners has been reported to be strong in many different mutualistic systems, such as pollination (34), seed-dispersal (32, 40), mycorrhizae (41) and cleaner-client fishes (42), we expect that these systems would exhibit a stabilizing seasonal structure. Our results could explain why even tropical mutualistic systems with relatively constant abiotic conditions, such as plant-hummingbird interactions, exhibit a stabilizing seasonal structure (11, 15).

The phenological asynchrony between species sharing partners has been used in several empirical systems to evidence phenological coevolution driven by competition for mutualistic partners (43–45). Our model provides theoretical support to the fact that phenological asynchrony between species sharing partners, showed by a high proportion of “V+” motifs, suggests a stabilizing seasonal structure shaped by competition for mutualistic partners. Using 17 empirical plant-pollinator interaction networks, we showed a clear over representation of “V+” motifs, relative to a random distribution of phenologies. This empirical result verifies our theoretical findings and suggests that phenological coevolution, driven by competition for mutualistic partners, can shape seasonal structure of natural systems, although these systems also exhibit abiotic forcing. Consistently with previous results on mutualistic community (5, 11) and food webs (46), we found that neglecting this seasonal structure (*i*.*e*. aggregating interactions temporally) led to an underestimation of their structural stability of ∼20%, providing further support to the fact that the temporal dimension of interaction networks is key to understand their stability (37, 47).

We also highlighted that phenological traits are key to understand how coevolution shapes network structure. Most of the previous work studying the consequences of trait coevolution for the structure of mutualistic networks have failed to reproduce the nested structure that is observed empirically (48, 49), but see *ref*. (19). However, these previous works have focused on the coevolution of morphological traits, not considering phenological traits. Here, we show that the coevolution of morphological and phenological traits lead to interaction networks with different structures. Consistently with the previous results (48, 49), we found that the coevolution of morphological traits led to a collection of sub-communities, each comprised of a few specialist species, corresponding to a modular network. In contrast, the coevolution of phenological traits led to a more permissive resource partitioning, with networks containing some generalist species and a nested structure where more specialized species interact with the core of generalist species. Our results thus suggest that phenological traits, which have been documented as a structuring trait in many mutualist systems – pollination (14, 15), seed-dispersal (50, 51), ant-plant (52), etc. – might be key to understand how nestedness can emerge from coevolution.

Our findings linking phenological coevolution and increased community stability can be generalized to other kinds of traits, provided that they allow to decouple the interactions with a common partner in time or space. For example, one can consider traits associated with thinner temporal dynamics, such as pollinator’s daytime activity and flower opening times (e.g. diurnal *vs*. night; 53, 54) or morphological traits with inter- or intra-individual variation, such as flower heights (55). These kinds of traits can also create “V+” motifs, maintaining mutualistic interaction with two different partners while removing direct competition between them. For instance, differences in flower heights in a tree could allow two pollinator species that fly at different heights to avoid competition while still interacting with the same tree population. This can happen both if different individuals display different heights, or within the same individual tree when it has flowers at different heights. This vertical stratification exists for example in plant-hummingbird interactions (56). Thus, the results obtained here, with phenological and morphological traits, could be generalized to traits with and without inter-individual variation, respectively, which is consistent with recent results highlighting intraspecific variability as a key mechanism promoting the stability of plant-pollinator communities (57).

By showing that complex interaction structures fostering stability can emerge from species’ fitness driven evolution, we effectively link coevolutionary mechanisms and community ecology. Most importantly, the emergence of these stabilizing structures depends on which kind of traits coevolve. For example, the well-studied nested structure emerged when including phenological traits but not when considering only morphological traits. Overall, our results suggest that coevolutionary dynamics enhance the positive effect of seasonal structures on community stability (5, 10, 11, 36, 37), which is probably a key mechanism to understand how natural communities can be both diverse and stable.

## Acknowledgements

We thank Nerea Montes for the help while collecting data. We also thank the taxonomists L. O. Aguado and T. Wood for their help with some bee identifications. FD was funded by *postdoc Mobility* grant from the SNF (project number: P500PB_217801). VDG was funded by a Marie Skłodowska-Curie grant (N°101064340) from the European Union’s Horizon 2021 research and Emergia grant agreement DGP EMEC 2023 00146 from Junta de Andalucía. Data collection was funded by a Marie-Curie Career Integration from the European Union’s Horizon 2020, the BeeFUN project (PCIG 14-GA-2013-631653), granted to IB.

## Data availability statement

Codes to perform simulation, to do figures and to analyse empirical networks are available here: https://github.com/f-duchenne/Evolution_pheno_vs_morpho

Empirical plant-pollinator interaction data are available here: https://doi.org/10.5281/zenodo.13270451

## Competing Interest Statement

The authors declare no competing interests.

## Author contributions

FD designed the study and performed all analyses with inputs from VDG and IB. FPM collected the data, with punctual help from IB, and identified plant and insect species. FD wrote the first draft and all authors participated to editing and writing the final version.

## Notes

https://doi.org/10.5281/zenodo.13270452

### Competing Interest Statement

The authors have declared no competing interest.

### Summary of Updates

We have changed the numerical implementation of the model to make our model consistent to what is usually done in quantitative genetics and corrected the introduction. Results and conclusion did not change.

