## Supplementary figures and Methods for "Coevolution of phenological traits shapes plant-pollinator coexistence"

### Coevolution and temporal dynamics of species interactions shape species coexistence

#### Supplementary Methods I

*Bounding the parameter values of the traits*

To avoid infinite evolution we mimicked physiological limits, we bounded the mean trait values between 80 and 285, and the standard deviation values from 1 to 50. To do so, we used a modified logit function for each parameter. The mean was transformed as follow:

$$\mu = \frac{80 + 205e^{\ddot{\mu}}}{1 + e^{\ddot{\mu}}} \qquad (S1)$$

The standard deviation was transformed as follow:

$$\sigma = \frac{1 + 49e^{\ddot{\sigma}}}{1 + e^{\ddot{\sigma}}} \qquad (S2)$$

*Equation for pollinators*

In the main text we only show the equation for the plant side, pollinators being described by similar equations. However, for clarity, here we present the similar equations describing competition terms and fitness of pollinators.

The competition between pollinators was described by the following equation.

$$\alpha_{jk} = \tau_{jk}^{A} \frac{\sum_{i=1}^{n_P} \gamma_{ij}\gamma_{ik}}{\frac{1}{2}\sum_{i=1}^{n_P} \gamma_{ij} + \gamma_{ik}} \qquad (S3)$$

where $n_P$ was the number of plant species and $\tau_{jk}^{A}$ the within guild phenological overlap between plant *i* and *k*.

The fitness function for pollinator *j* was:

$$W_j = -c\sum_{k=1}^{n_A} \alpha_{jk} + \sum_{i=1}^{n_P} \gamma_{ij} \qquad (S4)$$

#### Supplementary Methods II

*Network structure*

To assess how coevolutionary dynamics affect the structure of the interaction network, we measured indices describing the mutualistic interaction network. Since our interaction networks were fully connected but sometimes with links that were extremely weak, we rounded the interaction strengths ($\gamma_{ij}$) to the third digit. We measured the weighted connectance, nestedness, modularity, specialization and interaction evenness of the mutualistic networks. As different indices gave different information, we measured nestedness using two different indices, the index proposed by Galeano *et al.* (Galeano *et al.* 2009) and the wNODF (Almeida-Neto *et al.* 2008). All these indices were computed with the *bipartite* R package (Dormann *et al.* 2008). For the specialization, we used the $H_2$' index described in Blüthgen *et al.* (2006). The weighted connectance metric implemented in the *bipartite* R package is not sensitive to shifts in scale but only to the distribution of interactions. To track how coevolutionary dynamics was affecting the strength of mutualism we also calculated the average mutualism strength, which is the average of all $\gamma_{ij}$ and be view as a second metric of connectance.

Since evolution slowed down with time (Fig. S5), we measured these metrics at 10 time steps, that were evenly distributed on a square-rooted time scale (t = 0, 20, 100, 220, 400, 620, 890, 1210, 1580, 2000). To track network structure over time, we performed a Principal Component Analysis (PCA) with the seven metrics described above, across all simulations and time steps. We used the two first dimensions of this PCA which explained 86% of the total variance, to characterize the evolutionary trajectories of the simulated communities.

*Mutualism-competition balance and motifs promoting facilitation*

To assess how coevolutionary dynamics balance mutualism and competition at the community level, we also calculated the average competition strength, which the average of all $\alpha_{ij}$, and the balance between mutualism and competition as $\frac{\gamma_{ij}}{c\alpha_{ij}}$.

To test if co-evolutionary dynamics can lead to phenological motifs that promote facilitation over competition, we calculate the number of motifs with a “V” shape promoting facilitation, or “V+” motifs (Fig. 2e). This kind of motifs can be present only in the phenological scenario. In contrast, when considering morphological traits, all interactions happen simultaneously ($\tau = 1$) and thus competition between two species from the same guild does not depend on their trait overlap but only depends on their overlap in mutualistic interactions.

Since our in-silico mutualistic networks were weighted and without real zeros, we had to put a threshold above which we consider mutualistic interactions were strong enough to constitute “V” shape motifs. We used only the links for which $\gamma > 0.01$ and we extracted all corresponding “V” motifs. We calculated the number of “V+” motifs as those where the two species from the same guild overlapped less in their trait value than the product of their overlap with their mutualistic partner. For example, a “V” motif in which a plant $i$ interact with pollinators $j$ and $k$ would be “V+” motif if $\gamma_{ij}\gamma_{ik} > \tau_{jk}$, and “V-” motif otherwise (see Fig. 1). As connectance was changing among simulations because of different initial conditions, and within simulations, because of coevolutionary dynamics, we estimated the proportion of “V+” motifs, over the total number of motifs with a “V” shape. This allowed to get comparable measures over time and across simulations.

*Structural stability*

To understand how evolutionary dynamics affect the ecological stability of our system, we measured the community structural stability. Structural stability is a measure of ecological stability that is uses on generalized Lotka-Volterra models to quantify the probability of coexistence of a given species et, according to their interactions. Here, we defined the Lotka-Volterra model using the fitness function of each species to model their population dynamics. In this model, the variation in the population size of plant $i$ ($N_i$) and pollinator $j$ ($N_j$) were described by the following equations, respectively:

$$\frac{dN_i}{dt} = N_i\left(r_i - c\sum_{k=1}^{n_P}\alpha_{ik}N_k + \sum_{j=1}^{n_A}\gamma_{ij}N_j\right) \qquad (S5)$$

$$\frac{dN_j}{dt} = N_j\left(r_j - c\sum_{k=1}^{n_A} \alpha_{jk}N_k + \sum_{i=1}^{n_P} \gamma_{ij}N_i\right) \qquad (S6)$$

where $r_i$ was the basal per-capita growth rate of species $i$, and $c$ was the overall importance of competition, relative to mutualism, and similarly for pollinators.

Structural stability quantifies the probability of coexistence as the normalized range of environmental conditions, modelled through the per capita basal growth rates ($r$), in which all species can coexist in a stable way (Rohr *et al.* 2014; Saavedra *et al.* 2016; Song *et al.* 2018). The range of environmental conditions, or growth rates, in which all species can coexist is called the feasibility domain, and the structural stability is quantified as the size of the feasibility domain scaled by the number of species in the community, to be compared across communities with different number of species.

To calculate the structural stability, we needed to find all the $r$ values leading to positive abundances for all species. The vector of species abundances at equilibrium ($N^*$) was given by $N^* = A^{-1}r$, where $r$ was the vector of species growth rates and $A$ was the adjacency matrix describing within guild interaction (competition) as well as between guilds interactions (mutualism) among all pairs of species. This matrix was composed by four blocks:

$$A = \begin{bmatrix} -c\alpha_{(P)} & \gamma \\ \gamma^T & -c\alpha_{(A)} \end{bmatrix} \qquad (S7)$$

where $\alpha_{(P)}$ was the competition matrix among plants, dimensions of $n_P \times n_P$, and $\alpha_{(A)}$ was the competition matrix among pollinators ($n_A \times n_A$). $\gamma$ was the mutualistic interaction matrix ($n_P \times n_A$), and $\gamma^T$ its transposed.

The feasibility domain is the range of vectors $r$ that give $N^*$ with positive species' abundances only. The size of the feasibility domain ($\Omega(A)$), was calculated as follow:

$$\Omega(A) = \frac{vol\left(D_f(A) \cap B^S\right)}{vol(B^S)} \qquad (S8)$$

where $D_f(A)$ was the feasibility domain of $A$ and $B^S$ was the closed unit ball in dimension $S = n_P + n_A$.

To quantify structural stability, we normalized the size of the feasibility domain by the number of species in the community: $\omega(A) = \Omega(A)^{1/S}$. The normalized feasibility, $\omega(A)$, is bounded between 0 and 1 and corresponds to the average species persistence probability in the community, represented by the interaction matrix $A$. The closer the feasibility is to one, the more likely is that all species will coexist in a stable way. To compute this normalized feasibility, we used the *Omega* R function provided by Song *et al.* (Song *et al.* 2018).

*Competition structure and the structural stability*

The introduction of a competition for mutualistic partners within the competition terms ($\alpha$) of the generalized Lotka-Volterra model has been debated when studying mutualism, especially when using structural stability (Bascompte & Ferrera 2020; García-Callejas *et al.* 2023). Here we develop why we think the structure of the competition for mutualistic partners is important and should be included in the $\alpha$, in line with *ref.* (García-Callejas *et al.* 2023).

In linear generalized Lotka-Volterra model of bipartite interaction networks, such as our model, the fixed point can be written using effective competition and effective growth rate terms as follow (Bascompte & Ferrera 2020; Rohr *et al.* 2014), for plants:

$$R_i^{(P)} - \sum_k^{n_P} C_{ik}^{(P)} N_k^{(P)} = 0 \qquad (S9)$$

where $R_i^{(P)}$ is the effective growth rate of the focal plant species and $C^{(P)}$ refers to the effective competition matrix for plants. A similar equation describes the fixed point for pollinators, using $R_i^{(A)}$ and $C^{(A)}$, the effective growth rate of the focal animal species and the effective competition matrix for animals, respectively. When using linear functional response, effective competition and growth rates are given by:

$$R_i^{(P)} = r_i^{(P)} + \sum_j^{n_A} \sum_l^{n_A} \gamma_{ij} \left(\alpha_{jl}^{(A)}\right)^{-1} r_j^{(A)} \qquad (S10)$$

$$C_{ik}^{(P)} = \alpha_{ik}^{(P)} - \sum_j^{n_A} \sum_l^{n_A} \gamma_{ij} \left(\alpha_{jl}^{(A)}\right)^{-1} \gamma_{kl} \qquad (S11)$$

The term $\gamma_{il}\left(\alpha_{lj}^{(A)}\right)^{-1}\gamma_{kj}$ describes the effect of mutualistic interaction between the focal plant *i* and animal *j* corrected by competition between animals *l* and *j* enhanced by the mutualistic interaction between plant *k* and animal *j*. This emphasizes that effective competition integrates direct competition ($\alpha_{ik}^{(P)}$) but also indirect effects via mutualistic partners. This led previous work to conclude that in such models competition ($\alpha_{ik}^{(P)}$ and $\alpha_{ik}^{(A)}$) should include only competitive effects arising from limiting factors not explicitly present in the model (*e.g.* competition for water or light) and should not include competition for mutualistic partners (Bascompte & Ferrera 2020). We think that this is an erroneous interpretation of the effective competition. Effective competition account for indirect effects from plants to plants through the competition between pollinators ($P_k \rightarrow^{+} A_l \rightarrow^{-} A_j \rightarrow^{+} P_i$) but does not model direct competition arising from sharing partners, such as nectar depletion in pollination.

Here we decided to consider that direct competition was only due to interactions with shared mutualistic partners. We know that within each guild's species would compete for other limiting factors that were not in the model (*e.g.* competition for water or light for plants), but since here we were only interested by the evolutionary dynamics emerging from mutualistic interactions and their byproduct (i.e. competition for partners), we did the choice to ignore competition for non-modelled factors.

#### Supplementary Methods III

If evolution of phenologies drove, at least partially, the seasonal organization of the interactions, we expect to find a non-negligible proportion of "V+" motifs, in line with what we found in coevolved seasonal structures. To do so, we estimated species phenologies. Since sites were all in the same geographical region (maximal pairwise distance was 53km), we assumed that phenologies were the same in all sites and thus we used all the data available to infer phenological parameters, which allows to get more precise phenological estimates per species. We also assumed gaussian phenologies for all species, meaning that we had to estimate only two parameters, the mean and the standard deviation.

*Empirical flowering phenologies of plant species*

To calculate plant flowering phenologies we used the data of the flower count transects realized while collecting the plant-pollinator interactions data. We assumed gaussian phenologies, and thus had to estimate only two parameters: the mean and the standard deviation. We pooled all sites and years together, to gain power and get better estimates of the mean and standard deviation of the flowering dates. We estimated the mean flowering date as the average day of the year of flowering events observed, weighted by the number of flowers observed. Similarly, we calculated the standard deviation as the square root of the variance of the flowering dates, weighted by the number of flowers. To do so, we used the square root of the weighted variance, calculated using the *wtd.var* of the *Hmisc* R package (Harrell Jr 2023). For the plant species with one observation or with a null standard deviation, we set the standard deviation to one.

*Empirical activity phenologies of pollinator species*

To calculate pollinator activity phenologies we used the interaction data collected as we have no independent measure of their activity period. As for plants, we assumed gaussian phenologies and we pooled all sites and years together, to gain power and get better estimates of the mean and standard deviation of the activity periods. We estimated the mean activity date as the average day of the year of observations, weighted by the number of individuals observed. Similarly, we calculated the standard deviation as the square root of the variance of the flowering dates, weighted by the number of individuals. For the pollinator species with one observation or with a null standard deviation, we set the standard deviation to one.

*Proportion of V+ motifs in empirical networks*

Once we have estimated the species phenologies, we counted the number of "V+" motifs, in which the (geometric) average overlap between mutualistic partners was higher than the overlap between the two competitors. To get a measure comparable with our simulations and among networks, as before, we estimated the proportion of "V+" motifs over the total number of "V" shape motifs.

*Structural stability of empirical networks*

To calculate the structural stability of the empirical networks, we ideally want to get mutualistic interaction strengths that do not depend on sampling effort, neither on species abundances. To correct interaction strength by sampling effort, we calculated the average number of detected interactions per sample round: $\overline{I_{ij}} = \frac{1}{N}\sum_{s=1}^{N} I_{ijs}$, where *N* is the number of sampling events, and $I_{ijs}$ the number of interactions detected between plant *i* and pollinator *j* during sampling event *s*. To account for species abundances, we divided the average number of interactions by the geometric mean of the marginal sums of species involved in the focal interaction:

$$\gamma_{ij} = \frac{\overline{I_{ij}}}{\sqrt{\sum_{i=1}^{n_P} \overline{I_{ij}} \times \sum_{j=1}^{n_A} \overline{I_{ij}}}} \qquad (S12)$$

From the $\gamma_{ij}$ we could calculate the within guild competition terms $\alpha_{ik}$ using the equation (3) of the main text. As in simulations, to neglect the seasonal structure in interactions we set $\tau_{ik} = 1$, and to include the seasonal structure in interactions we calculated $\tau_{ik}$ according to equation (2), using the empirical phenologies. However, since for some networks interspecific competition reached one, we fixed the intraspecific competition to $\alpha_{i,k=i} = 1.05$ to insure diagonal dominance.

Finally, to see how considering the seasonal structure in interactions affected the structural stability of these communities, we estimated the feasibility ($\omega$) with and without considering the seasonal structure in the interactions. We used the generalized linear Lotka-Volterra from equation (4) of the main text to calculate the structural stability of the community, with and without seasonal structure in interactions in the same way as described previously, for the different values of competition importance ($c$) that we used in the simulations (2, 4 and 6).

#### Supplementary Figures

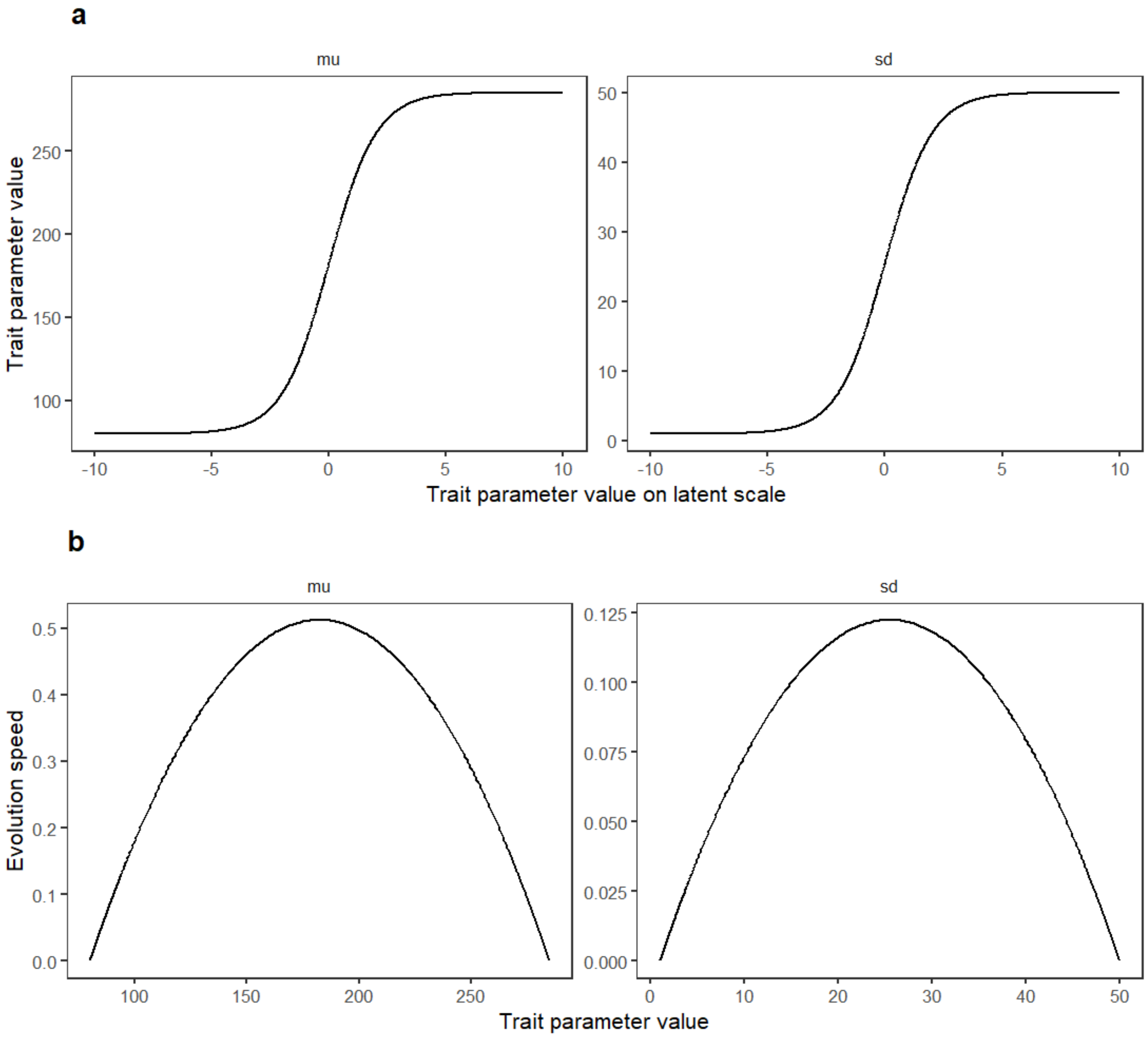

***Figure S1: Latent scale of trait parameters and evolution speed.*** *(a) Relationship between real values of the trait parameters and their value on the latent unbounded scale, for both parameters, the mean (μ, mu) and the standard deviation (σ, sd). (b) Evolution speed, measured as the additive genetic variance (g) once back-transformed to the real parameter value scale, as a function of trait parameter value, for both parameters. Since selection was applied on the latent unbounded scale, its effect on the trait value decreased gradually towards zero when approaching bounds of the parameter intervals, which allowed to keep trait parameter values in the chosen interval.*

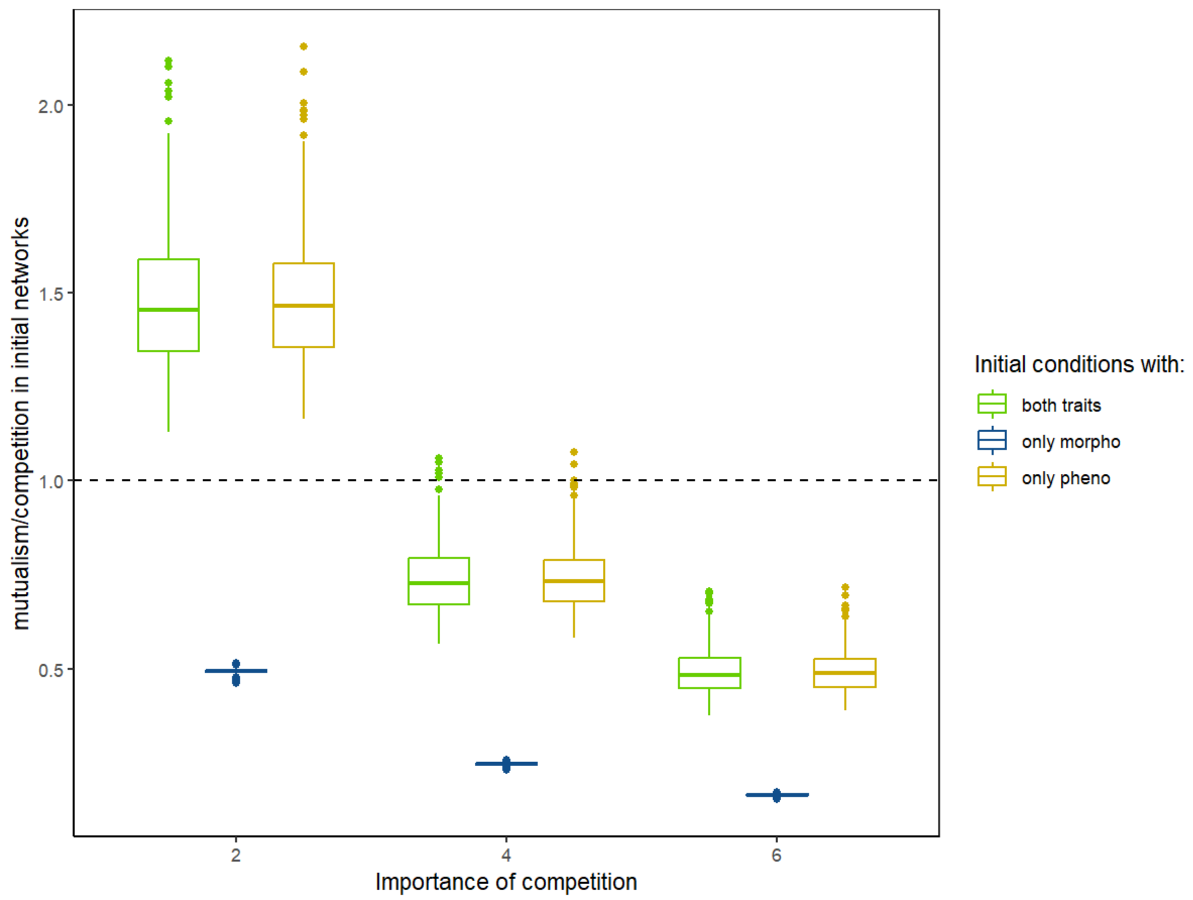

***Figure S2: Choice of values of competition importance.*** *Balance between mutualism and competition ($\frac{\bar{\gamma}}{c\bar{\alpha}}$, cf. methods of the main text) as a function of competition importance for the 100 initial conditions. The three values of competition importance (c) were chosen to have an average ratio of mutualism/competition across the 100 initial conditions when considering both traits that was above one when competition had low importance, slightly below one when competition was moderate, and below 0.5 when competition had strong importance.*

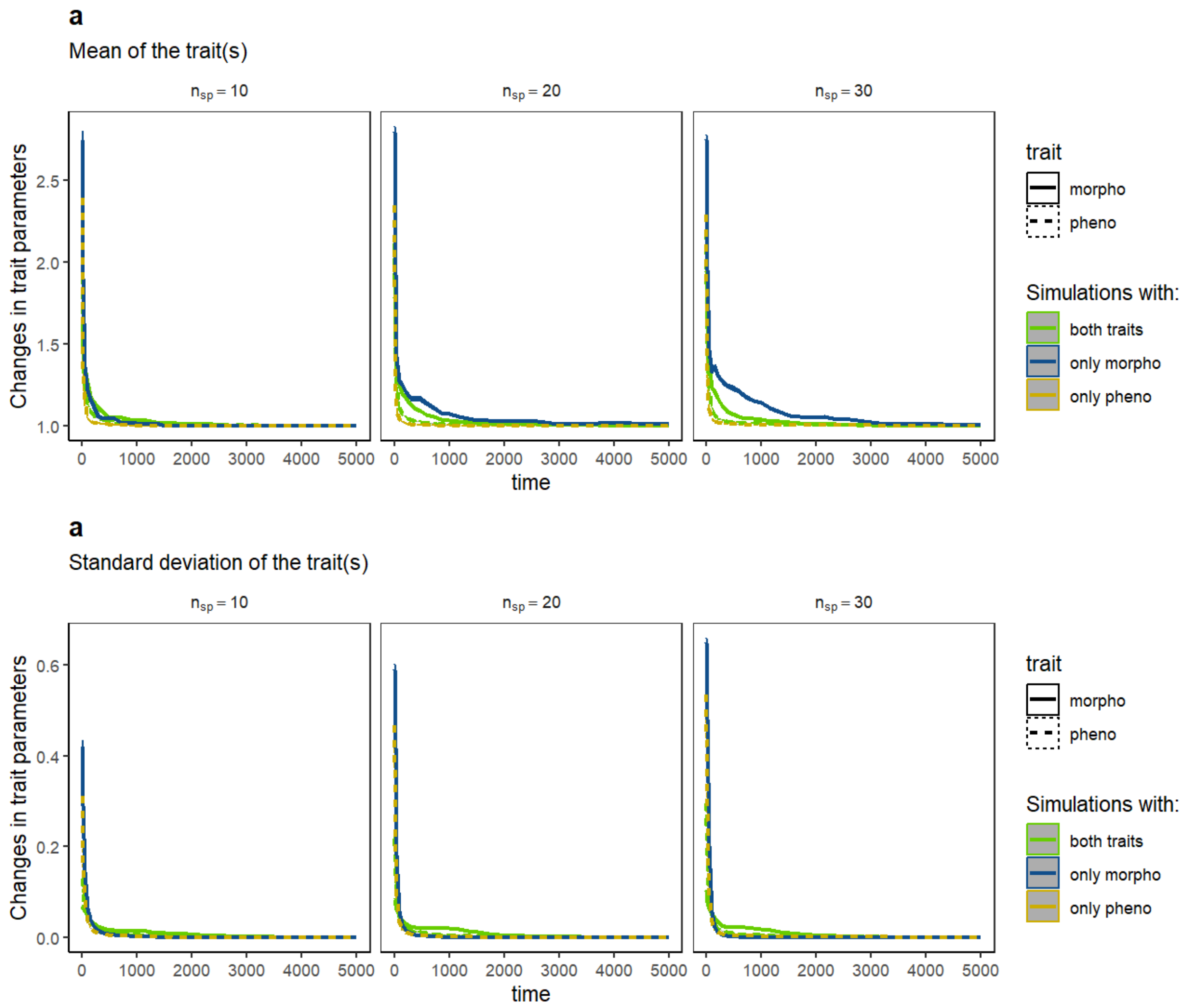

***Figure S3: Speed of trait evolution over simulations.*** *Temporal changes in traits parameters (per time step), mean (mu) and standard deviation (sd) on the modified logit scale (cf. Methods). The speed of trait evolution drop over time to reach very low values. The line and ribbon (too small to be visible) represent the mean and the associated 95% confidence interval, respectively. Changes in trait parameters were average across all species and across the 100 simulations for each number of species per guild (*$n_{sp}$*). Only the simulation performed with a strength of competition c = 4 are shown here.*

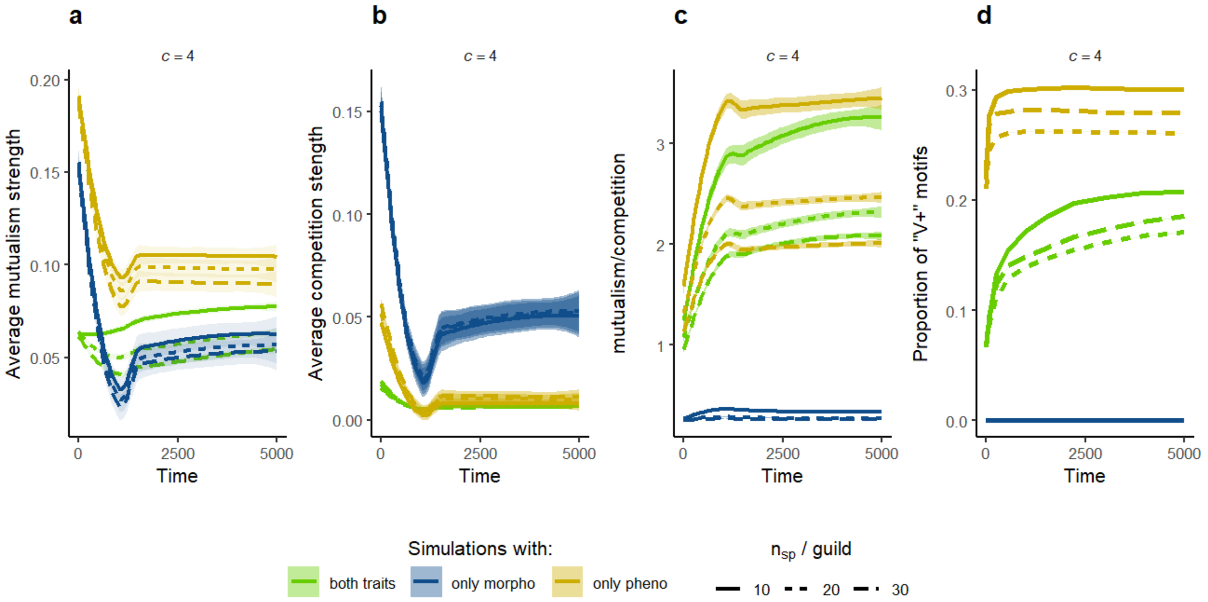

***Fig. S4: motifs promoting facilitation emerge from phenological coevolution.*** *Same as Figure 2 of the main text but for c= 4. (a) average mutualistic strength, (b) average competition strength, (c) mutualism/competition balance (cf. methods) and (d) proportion of "V+" motifs (over all "V" motifs), as a function of time, diversity. In (a)-(d) the lines and ribbon show the estimated values and their 95% confidence interval, respectively, obtained by a locally estimated scatterplot smoothing (LOESS) of the values across time. V+ motifs did not exist when performing coevolution with morphological trait only (cf. Methods).*

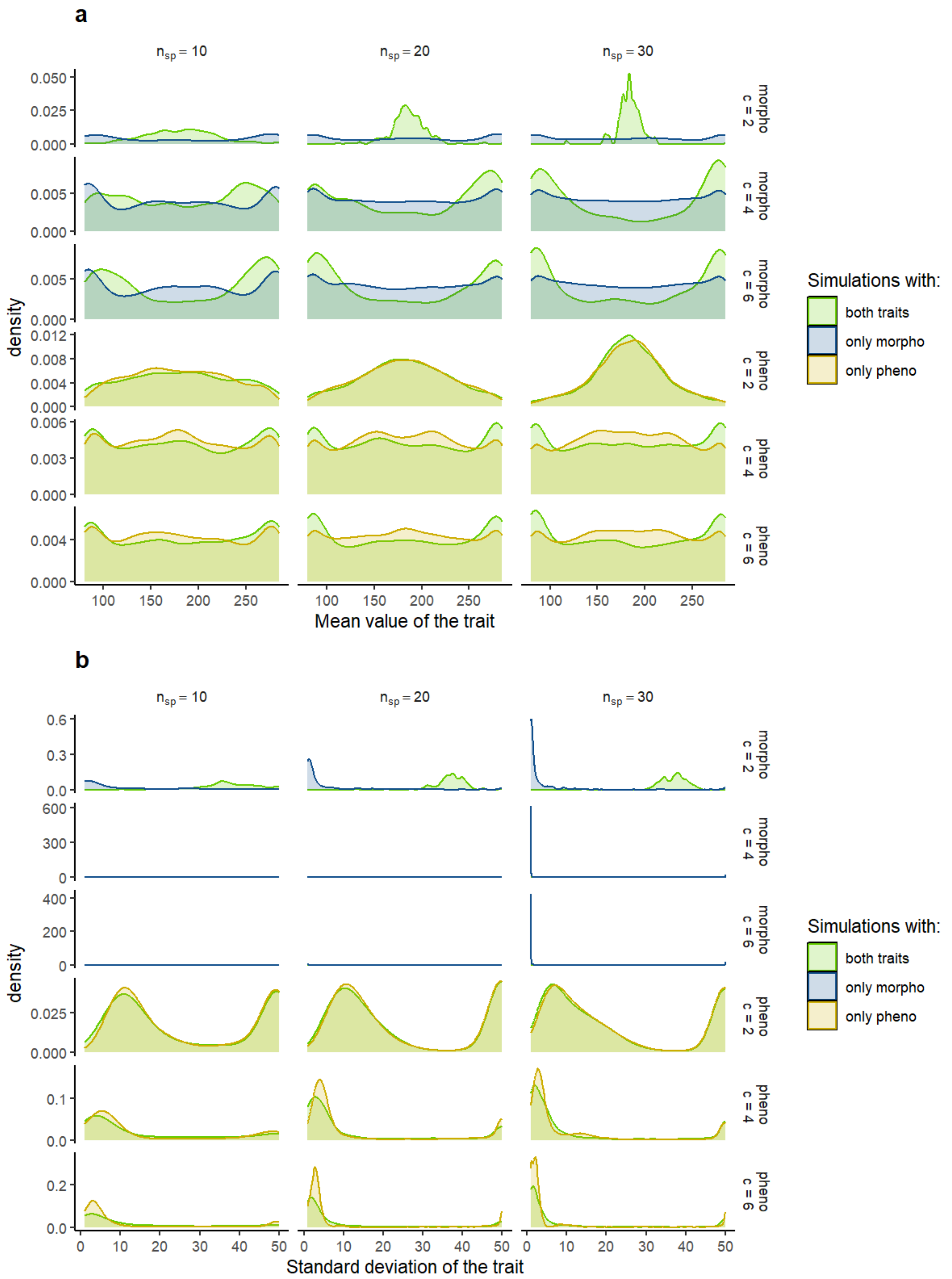

***Figure S5: Trait distribution at the end of the simulations.*** *Density distribution of (a) the mean trait values and (b) of the standard deviation of the traits at the end of simulation, as a function of the diversity (number of species per guild) and of the simulation scenario. Simulations done with an overall strength of competition c = 4. Morphological coevolution led to a bimodal or trimodal mean trait value distributions, with specialist species only (no large standard deviation), which correspond to a modular structure of the interaction network.*

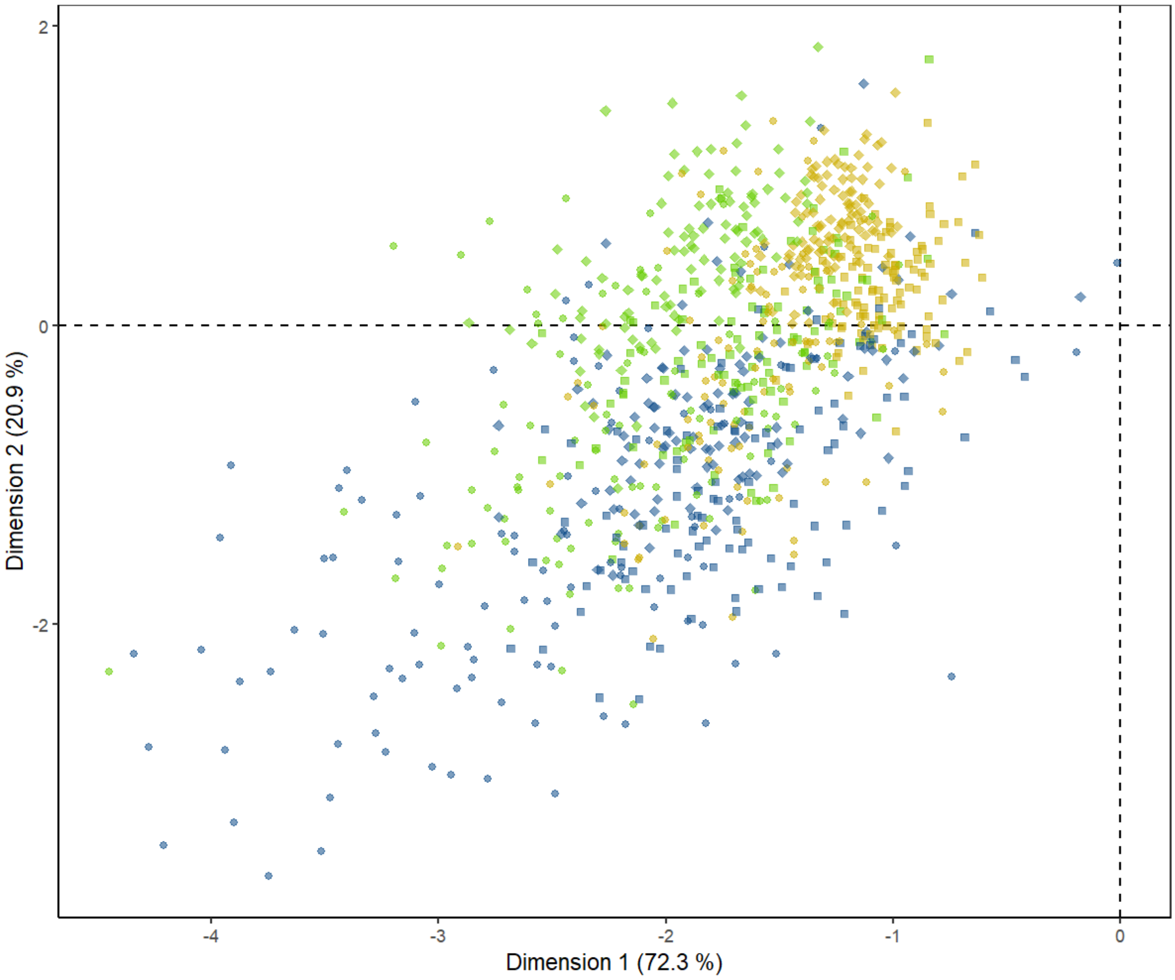

***Figure S6: divergent evolutionary trajectories of communities structured by phenological and morphological traits with low competition.*** *Similar figure as Fig. 3c of the main text, but when performing simulations with a lower strength of competition, c = 6 instead of c = 4. Circles represent simulations with 10 species per guild, squares 20 species per guild and diamonds 30 species per guild.*

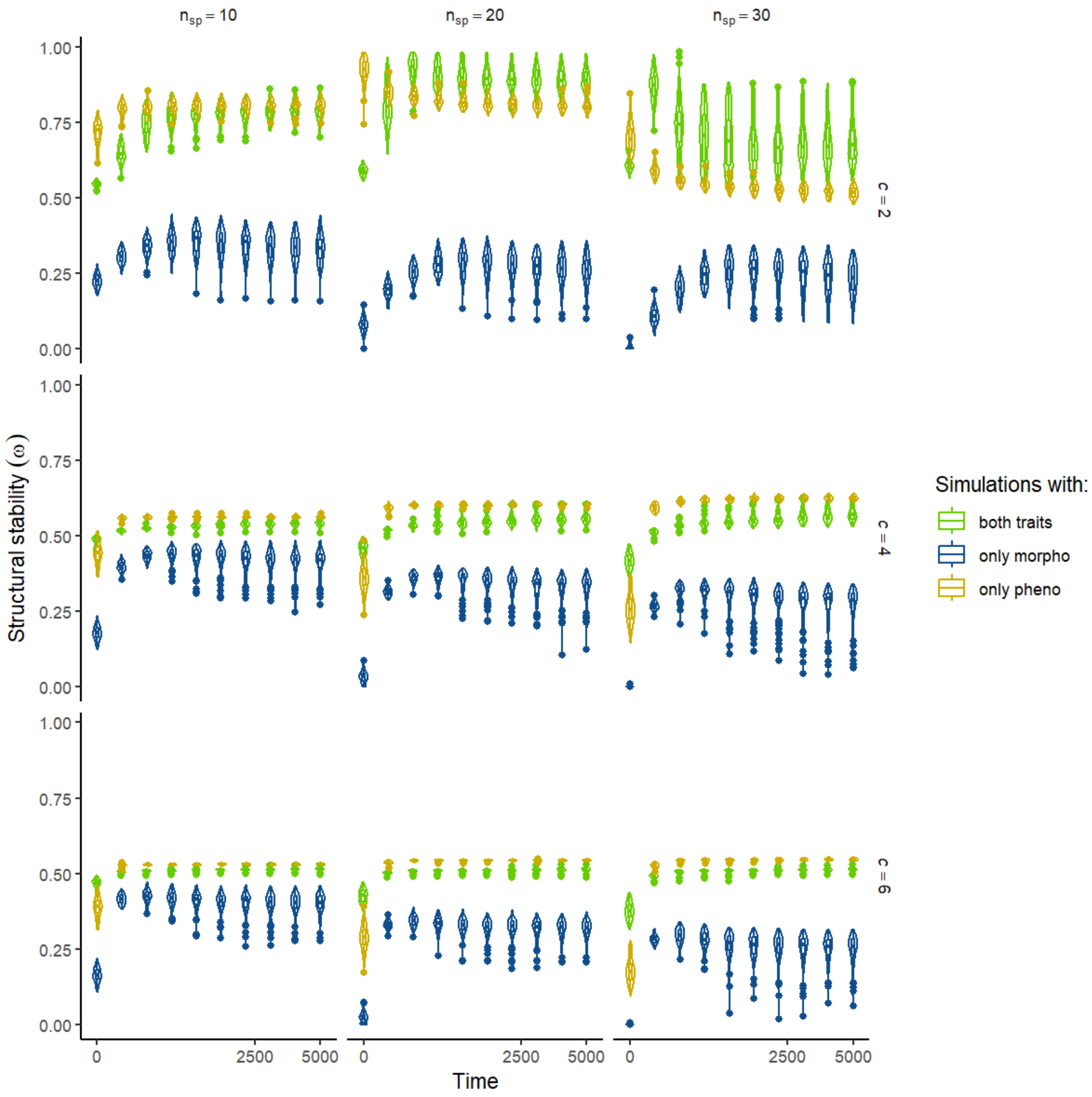

***Figure S7: effect of coevolution on structural stability when competition had low and high importance in the model.*** *Similar to Figure 3 of the main text but adding simulations with c = 2 and c=6. Changes in structural stability (size of the feasibility domain) over time, for each simulation scenario and number of species per guild. The x-axis is square-root transformed.*

#### Supplementary Table

**Table S1:** empirical interaction networks, their diversity and the sampling effort.

| Site | Number of pollinator species | Number of plant species | Total number of sampling events | Number of sampled years |
|---|---|---|---|---|
| Aznalcazar | 124 | 29 | 69 | 8 |
| Bonares | 103 | 24 | 63 | 8 |
| Convento_de_la_luz | 123 | 25 | 59 | 8 |
| Esparragal | 61 | 24 | 54 | 8 |
| La_cunya | 78 | 27 | 56 | 8 |
| La_rocina | 131 | 31 | 66 | 8 |
| Las_mulas | 57 | 18 | 22 | 3 |
| Niebla | 95 | 28 | 35 | 5 |
| Pinares_de_hinojos | 77 | 36 | 54 | 8 |
| Pino_del_cuervo | 111 | 32 | 63 | 8 |
| Urbanizaciones | 116 | 26 | 69 | 8 |
| Villamanrique_este | 108 | 26 | 64 | 8 |
| Villamanrique_sur | 84 | 20 | 60 | 8 |
| Cotito_de_santa_teresa | 75 | 22 | 34 | 5 |
| El_pinar | 21 | 8 | 6 | 1 |
| El_hongo | 53 | 14 | 24 | 4 |
| Laguna_del_ojillo | 57 | 17 | 21 | 2 |
